# QSProteome: A Community-Driven Interactive Platform for Large-Scale Exploration and Evaluation of Predicted Protein Complex Structures

**DOI:** 10.1101/2025.09.10.675416

**Authors:** Edward A. Catoiu, Delina Kambo, Brianna Rodriguez, Krista Vear, Ziheng Wang, June Akpata, Jaladhi Shah, Rayyan Ayoub, Matthew Hofmann, Keerthana Ananda, Avani Tantry, Ishita Gupta, Robin Anwyl, Carter Faucher, Jordan Nichols, Brian Ton, Lorenzo O. Marchal, Bernhard O. Palsson

## Abstract

QSProteome (https://QSProteome.org) is a community-scale platform for modeling, evaluating, and refining quaternary protein structure. The resource hosts 35,528 unique modeled assemblies spanning over 42,000 genes, covering nearly all curated complexes in BioCyc and ComplexPortal databases. Each model is displayed on an interactive page with 3D visualization, chain-level confidence metrics, structural alignments, functional annotations, and automated stoichiometry checks against database expectations. A cloud-based server supports continuous user uploads and automated processing pipelines—enabling submission and validation of >54,000 models within 14 weeks. To promote iterative refinement, QSProteome includes a gamified re-curation workflow that transitions users from training modules into live curation, enabling the community-led assessment of 1,547 ABC transporter complexes. Together, these components form a dynamic, scalable infrastructure for proteome-scale structural biology. By unifying modeling, validation, and annotation in a reusable, searchable, and community-extensible framework, QSProteome enables proteome-scale structure accessibility and reuse—powering discovery, annotation, and collaborative refinement across the structural biology community.

## Introduction

Recent advances in structure prediction—especially AlphaFold^1^ and ColabFold^2^—have enabled high-accuracy modeling of monomeric and oligomeric proteins, reducing reliance on experimental methods such as X-ray crystallography and cryo-EM^3^. AlphaFold’s performance in CASP14 demonstrated near-experimental accuracy for monomeric proteins^4^, and its multimer extension, AlphaFold-Multimer^5^, extended predictive capabilities to protein complexes, accelerating structural exploration across thousands of proteomes.

However, while modeling capabilities have advanced rapidly, the infrastructure for organizing, evaluating, and interacting with predicted oligomeric complexes has not kept pace. Oligomeric structure predictions are often released as static coordinate files buried in the supplemental data of accompanying publications, or dispersed across isolated repositories, lacking annotations, metadata, validation metrics, and links to biological context. ModelArchive^6^ offers archival support for computational models but provides limited interactivity or biological context. AlphaFoldDB^7^ highlights the value of interactive centralized structure repositories but lacks support for quaternary assemblies, leaving many predicted complexes scattered, unvalidated, and underutilized.

Moreover, the biological relevance of predicted quaternary structures remains difficult to evaluate without dedicated tools. While AlphaFold provides internal confidence scores—such as the inter-protein TM-score (ipTM), predicted TM-score (pTM), predicted Local Distance Difference Test (pLDDT), and Predicted Aligned Error (PAE)—these metrics do not always reflect the physiological plausibility of multimeric assemblies. Complementary validation approaches are needed to distinguish between biologically relevant complexes and non-physiological assemblies.

Developed independently of AlphaFold, the QSalign framework assesses evolutionary conservation of quaternary shape as a proxy for biological relevance^8-9^. By comparing predicted models to experimentally determined assemblies with conserved multimeric geometries, QSalign offers an orthogonal measure of structural plausibility^10^. Building on this approach, recent modeling efforts used AlphaFold 2 to predict thousands of homo-dimeric structures across four species—*E. coli, H. sapiens, P. furiosus, and S. cerevisiae*—validating them with QSalign and assembling higher-order oligomers using PISA, ultimately revealing that 20–45% of proteomes form homomeric assemblies^11-12^. However, these models were released as static zip files without annotation or validation, limiting their accessibility and utility.

To address this gap, we developed the Quaternary Structural Proteome Repository for Organized and Transparent Evaluation Of Modeling Efforts (QSProteome)—an open-access, interactive platform that supports community-wide organization, annotation, and evaluation of oligomeric protein structures. QSProteome transforms static structure predictions into interpretable and reusable resources. Each model page includes interactive 3D visualization, AlphaFold confidence metrics, structural alignments, automated quaternary shape (QS) validation and functional annotations from UniProt^13^, ComplexPortal^14^ and STRING^15^.

QSProteome enhances transparency in model evaluation by integrating quality metrics and comparative validation alongside biological context. To further support reproducibility and re-modeling efforts, the platform provides downloadable configuration files and AlphaFold 3-compatible input formats^16^.

By consolidating structural validation, confidence scoring, and biological annotation, QSProteome enables researchers to answer five fundamental questions for every predicted protein complex: (1) What does the protein structure look like? (2) Are the internal confidence metrics (pLDDT, pTM, ipTM, PAE and PAE-derived scores) consistent with a reliable model? (3) Is the quaternary shape of the modeled protein conserved in related structures? (4) Is there supporting evidence that the subunits form a physical complex? (5) What is the biological function of the modeled protein? This information–previously siloed across disparate tools and databases and accessible only through laborious case-by-case analysis–is now unified in a single interactive platform, enabling real-time access at unprecedented scale across tens of thousands of complexes and hundreds of species.

In addition to hosting and contextualizing models, QSProteome operates as a community modeling engine. Curated gene stoichiometries are extracted from protein complexes annotated in ComplexPortal and BioCyc (Tier 1 & 2 databases)^17-19^ and stored as “pending jobs”. Users can retrieve preformatted input files, model the structures, and upload results via a dedicated portal. Each job is uniquely assigned and tracked through a server-side queuing system (described below) to ensure non-redundant contributions. Upon submission, automated pipelines validate the model and enrich it with structural and biological annotations, enabling a continuous, crowd-sourced expansion of modeled proteomes.

Launched in April 2025, QSProteome hosts over 35,500 unique oligomeric models spanning 42,375 genes, supported by more than 157,000 structural alignments and nearly 8,500 quaternary shape evaluations. The resource is growing by approximately ∼400 models per day, driven by both automated batch uploads and community contributions. To sustain this rapid growth, its backend server is deployed on Amazon Web Services (AWS), using Amazon S3 (Simple Storage Service) for large-scale object storage, Amazon RDS (Relational Database Service) for structured data management, and Amazon EC2 (Elastic Compute Cloud) for on-demand processing and validation. This scalable cloud-based infrastructure, combined with integrated structure validation and functional context, positions QSProteome as foundational infrastructure for community-driven, structure-guided proteome modeling.

As a proof of concept for QSProteome, we targeted every ABC transporter annotated in BioCyc (Tier 1 and 2 databases), ComplexPortal, and previous large-scale modeling efforts^12,20-21^. We first generated AlphaFold 3 models using annotated subunit stoichiometries, then compared them against known quaternary architectures to flag misannotations. To crowdsource this validation, we launched the interactive ‘ABC Game’, which trains and certifies users to spot missing domains in 3D models. Ten certified players collectively reviewed 1,547 models (Table S1); those flagged were remodeled with corrected stoichiometries drawn from literature and homologous assemblies. This gamified workflow not only improved ABC transporter annotations and yielded higher-quality protein models, but also established a scalable, community-driven pipeline that is generalizable to the systematic refinement of other oligomeric protein assemblies.

## Database Description and Features

### 1. From upload to insight: automated processing of user-submitted AlphaFold models

The ability for any user to contribute models and receive proper attribution is central to QSProteome’s mission of building a centralized structural proteome repository. To support this, we provide an intuitive upload portal (Fig. 1a) where users can submit zipped AlphaFold 3 outputs^16^ (ColabFold^2^ models are supported on a legacy portal). QSProteome verifies required files, tracks upload status, and optionally collects attribution details.

**Figure 1.**
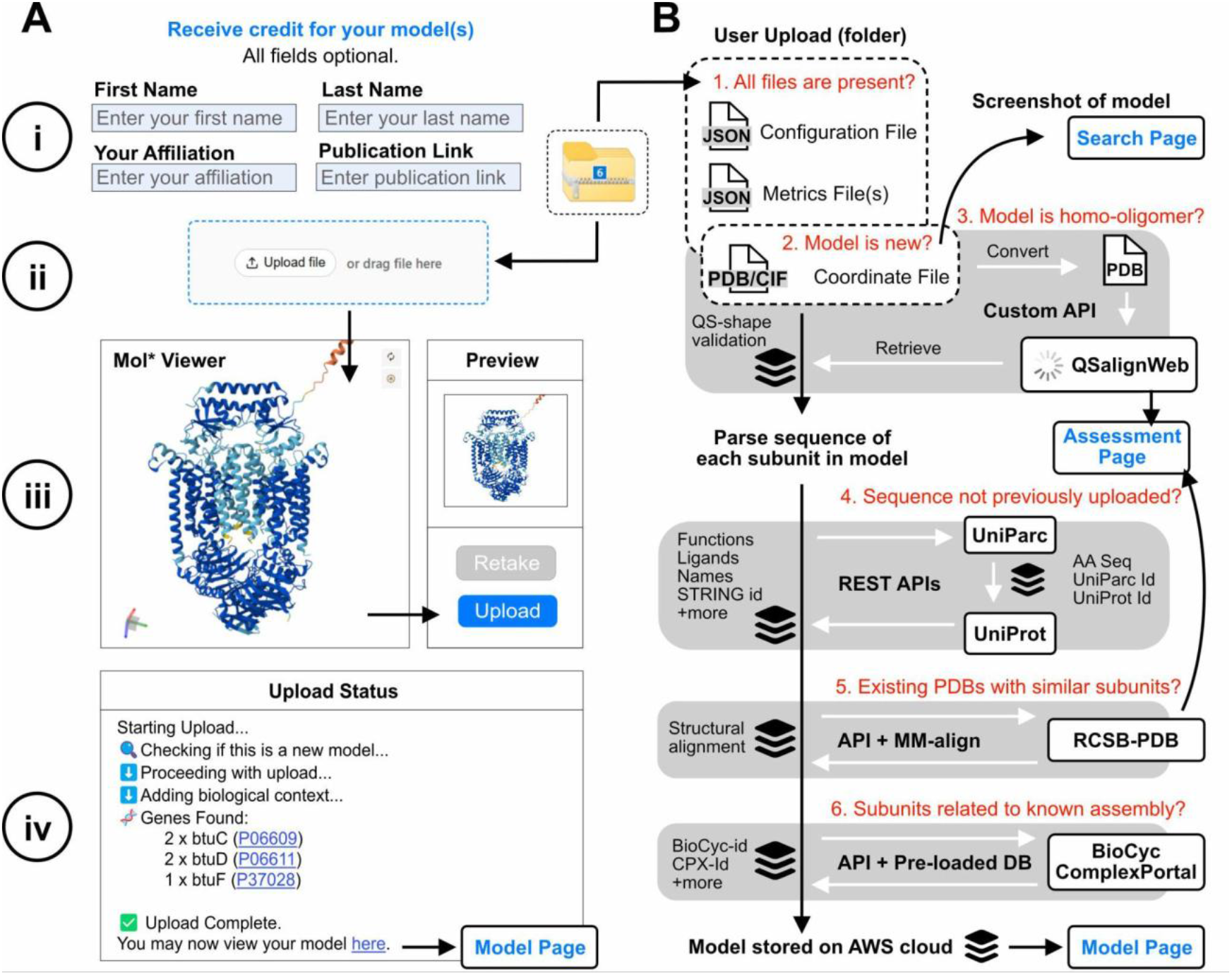
The QSProteome Upload Portal. **(A)** guides users through: (i) entering optional personal details for attribution, (ii) uploading a zip archive of up to 30 models, (iii) previewing each model, screenshotting (for search result rendering) and uploading, and (iv) tracking the status of uploaded models. **(B)** QSProteome automatically: (1) detects configuration, metrics, and coordinate files, (2) checks for duplicate uploads, (3) validates homo-oligomeric models via QSalignWeb, (4) retrieves sequence- and gene-level data from UniParc/UniProt^22^, (5) conducts structural alignments^24^ between the model and related structures in RCSB PDB^23^, and (6) maps models to external database entries. A screenshot of each model is stored for display on the Search Page. QSalign and MMalign aggregate results are displayed on the Assessment Page. Retrieved data is cached in server-side databases to minimize future queries to third-party APIs.

For each submission, the best-ranked model is selected and processed through an automated pipeline (Fig. 1b). To prevent duplication, coordinate files are verified using SHA-256 hash checks against previously uploaded models. Homo-oligomers are submitted to QSalignWeb for quaternary shape validation^10^. UniParc and UniProt annotations are retrieved for all subunits^22^, models are structurally aligned to related PDBs^23^ using MM-align^24^, and known assemblies are mapped from ComplexPortal and BioCyc. Each model is assigned protein- and gene-level metadata, a structure screenshot, and a dedicated webpage featuring 3D visualization, confidence scores, STRING interactions, and downloadable input files for AlphaFold 3.0. All annotations are retrieved via real-time API calls to ensure the displayed information remains current.

### 2. A cross-species structural repository with advanced search and filtering

As of this publication, QSProteome comprises *35,528 unique oligomeric models* spanning 222 NCBI taxonomy identifiers^25^, collapsed into 153 species-level taxa, including key model organisms (Fig. 2a). Generated with ColabFold (AlphaFold v2) and AlphaFoldServer (v3), these models capture the annotated gene stoichiometry of protein complexes curated in 84 BioCyc databases, ComplexPortal, and prior large-scale modeling efforts – including the *E. coli* quaternary structural proteome^20^, a whole-cell model of *Mycoplasma genitalium*^21^, and predicted homo-oligomeric assemblies across four diverse proteomes^12^. QSProteome hosts a model with an exact gene stoichiometric match for 94.9% of protein complexes in Biocyc and 99.9% in ComplexPortal (Fig. 2b). Modeling progress and quality metrics by species or dataset are summarized on the Repository page (qsproteome.org/repository) (Fig. 2c). For each dataset, only the highest-ranked model for each unique gene stoichiometry is displayed. Ranking scores have been previously defined for models generated with Alphafold 2 (see “Model confidence” of Evans et al 2022^5^) and Alphafold 3 (see Supplemental Methods 5.9.3 of Abramson et al., 2024^16^). Models with low internal confidence metrics (e.g., ipTM) may indicate assemblies whose gene stoichiometry warrants curation or refinement (see Fig. 6 and Fig. S1).

**Figure 2.**
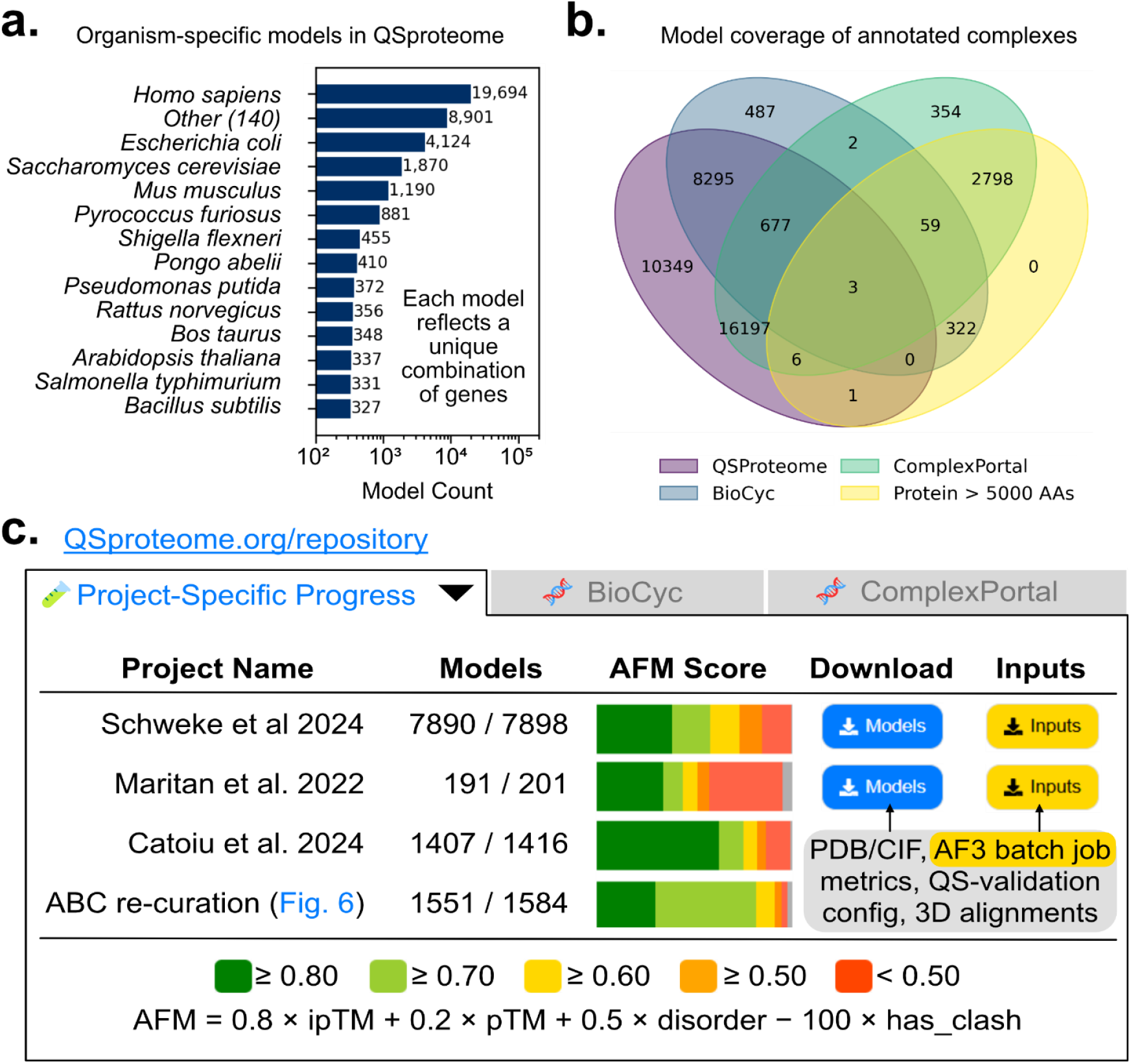
The size of the QSProtome Repository. **(A)** QSproteome hosts 35,528 unique structural models across 153 species-level taxa. Each unique gene-stoichiometry combination is shown once. **(B)** Gene signature overlap between QSproteome and annotated protein complexes. Proteins larger than 5000 amino acids or containing ambiguous sequence characters fail to produce a valid Alphafold 3 structure. **(C)** The Repository page summarizes modeling progress, displaying completed vs. target counts and color-coded AFM (ranking) scores (see Supplemental Methods 5.9.3 of Abramson et al., 2024^16^). Users can bulk-download pre-formatted AlphafoldServer inputs, models, and metadata to streamline batch submission. New projects encourage community participation. Only highest scoring unique models are shown. *Disorder* and *has_clash* are not used in AFM score calculation for models generated with Alphafold v2 (scoring metric defined by Evans et al., 2022^5^).

QSProteome features a flexible search engine that allows users to efficiently locate models using a wide range of identifiers. The primary search bar—available site-wide—supports keywords for UniProt IDs, gene names, synonyms, ligand names, ComplexPortal IDs, protein names, and functions (Fig. 3a). For structured queries, the Advanced Search allows filtering by organism, model availability, required genes in stoichiometry, confidence scores, total number of genes, and oligomerization type (Fig. 3b). Results display essential model attributes and can be sorted by confidence scores or complex size (Fig. 3c), with export options in CSV or JSON formats to support downstream analysis (Fig. 3d).

**Figure 3.**
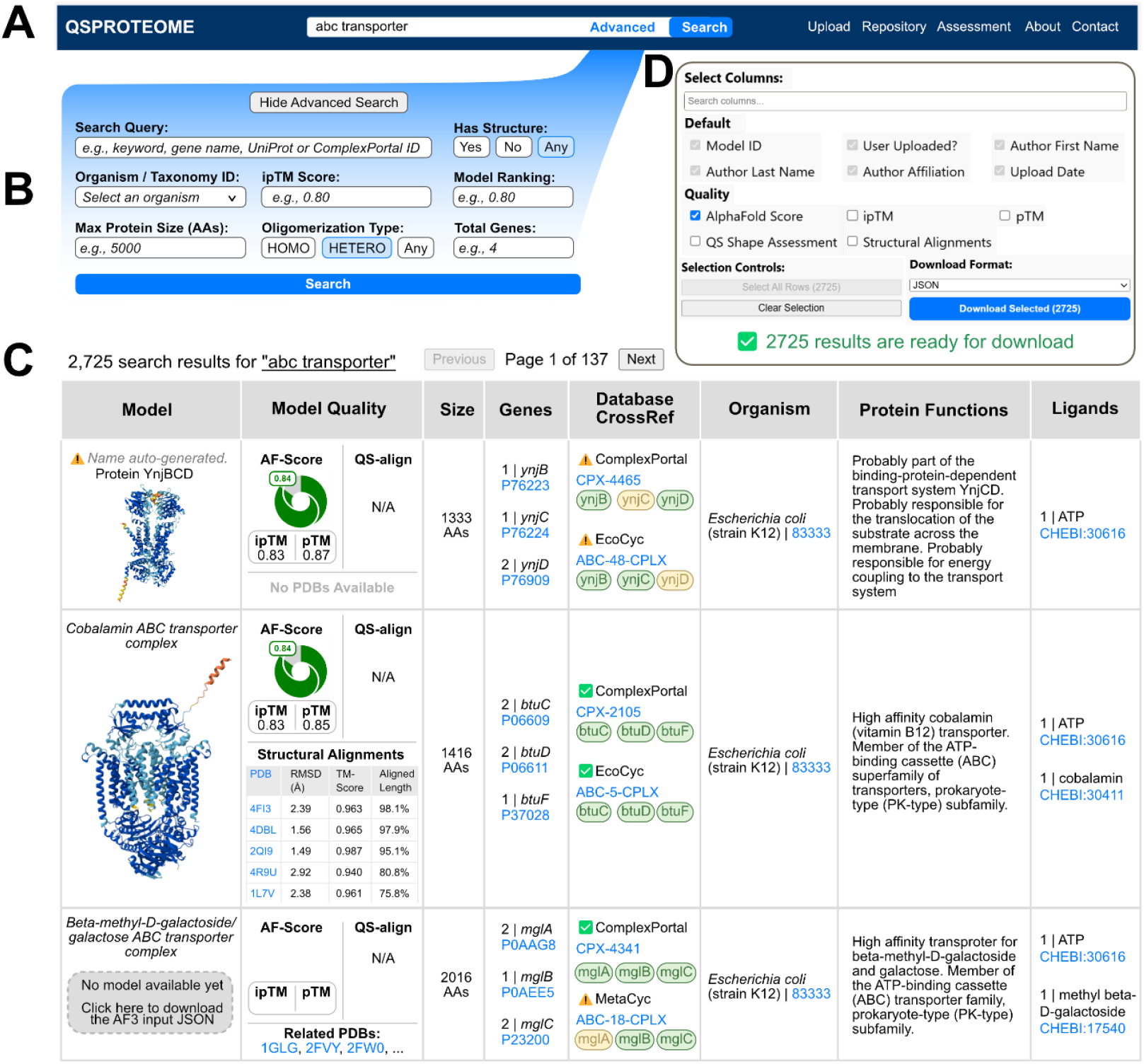
QSProteome Search and Retrieval Interface. **(A)** The keyword search retrieves models or known assemblies using gene names, protein IDs, functions, and ligands. **(B)** Advanced Search enables refined filtering by organism, gene name, model availability, complex size, confidence scoring, and oligomerization type. **(C)** Each result includes an image of the structure, AlphaFold metrics, QSalign validation (if available), MM-align alignments, complex size, gene stoichiometry, cross-referenced annotations, and ligands. Protein images link to dedicated model pages. All linked IDs (UniProt, ChEBI^26^, ComplexPortal, BioCyc, PubMed) redirect externally. Colored badges indicate stoichiometric matches; e.g., CPX-4465 includes an extra copy of *ynjC*. For unmatched complexes, downloadable AlphaFold 3 input files are available. **(D)** Results are exportable in JSON and CSV formats.

By combining intuitive keyword search with structured filtering and sorting, QSProteome enables researchers—specialists and non-specialists alike—to efficiently retrieve, analyze, and refine structural models, transforming predicted complexes from static archives into actionable biological resources.

### 3. A comprehensive framework for assessing model quality

To promote transparency and build confidence in computational models, QSProteome provides a robust evaluation framework with direct access to structural files, modeling parameters, and quality metrics—all presented in an intuitive, interactive format.

Each model page features an embedded Mol* viewer^27^ for real-time 3D structure exploration (Fig. 4a). Global confidence scores—including inter-residue predicted TM-score (ipTM), predicted TM-score (pTM), and a summary model confidence score—are automatically parsed to provide an overview of overall model reliability. Residue-level pLDDT scores and predicted aligned error (PAE) matrices offer finer resolution for assessing local accuracy and domain positioning (Fig. 4b). Newer, PAE-derived metrics for improved evaluation of interacting chains—e.g., chain-minimum PAE^16^ and ipSAE^28^ (interaction prediction Score from Aligned Errors) scores—are also displayed, demonstrating that QSProteome’s modular design enables emerging scoring metrics to be deployed across tens of thousands of models.

**Figure 4.**
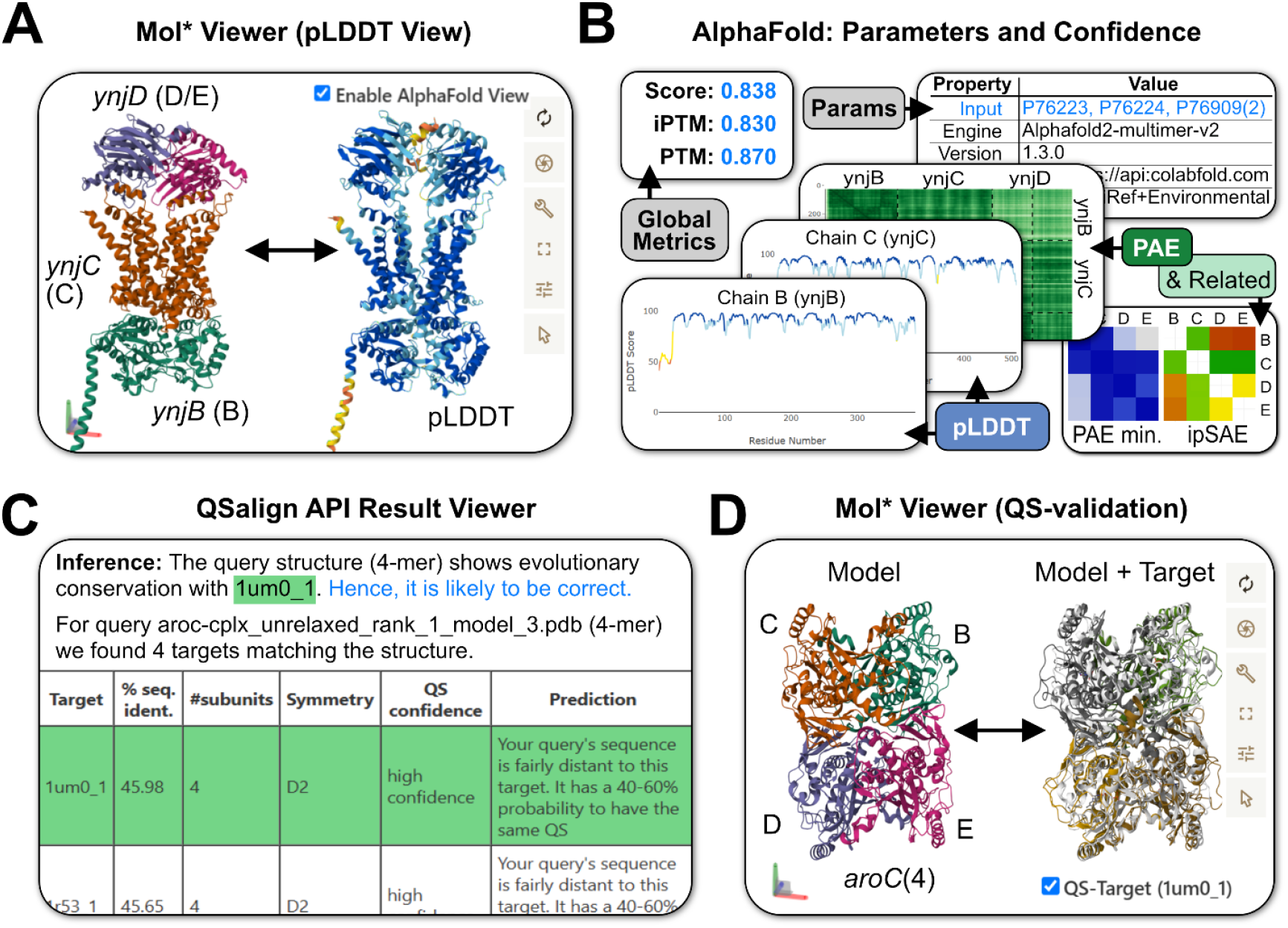
Transparent Model Assessment and Validation. **(A)** The embedded Mol* viewer allows interactive structure visualization with both standard and pLDDT-based color schemes. **(B)** AlphaFold metrics (ipTM, pTM, pLDDT, PAE, chain min PAE and ipSAE) are displayed alongside modeling parameters. **(C)** QSalign validation results for homo-oligomers report quaternary structure conservation, symmetry class, subunit count, and confidence scores. **(D)** Users can overlay their model with a QSalign-validated target to visually assess structural similarity. Similar functionality is enabled for related PDB entries (not shown).

To ensure reproducibility, QSProteome displays complete metadata for each prediction, detailing template usage, MSA strategy, recycle counts, and ensemble settings used during model generation. This enables users to trace modeling choices, compare workflows, and interpret confidence scores in context.

For homo-oligomeric models, QSProteome integrates QSalign^10^, a structural alignment tool that evaluates quaternary structure conservation by comparing predicted assemblies to experimentally determined homologs. Summary tables report sequence identity, subunit counts, symmetry classification, and structural confidence (Fig. 4c). Where applicable, the closest validated structure can be overlaid with the predicted model in the Mol* viewer for side-by-side comparison (Fig. 4d).

In parallel, MM-align^24^ is used to compare each model against all PDB structures containing shared subunits. These alignments are summarized on each model’s page, and aggregate metrics are provided on the Assessment Page (Fig. S2).

By integrating visualization tools, residue- and domain-level confidence metrics, and full modeling metadata, QSProteome allows users to critically assess each model’s reliability before applying it to large-scale structural or functional studies. This transparent evaluation system empowers users to assess model reliability for downstream applications, enhancing confidence in structure-based predictions.

### 4. Enriching structural models with biological context

Interpreting predicted structures requires more than 3D coordinates—biological relevance hinges on understanding function, interactions, and known complex formation. QSProteome integrates data from UniProt, ComplexPortal, and STRING to provide functional context for every uploaded model^13-15^.

At the sequence level, QSProteome retrieves UniProt annotations for each modeled subunit, including gene names, protein function, ligand interactions, and cofactors^13,22^. This information is particularly useful for homo-oligomeric proteins, whose oligomerization state is often unknown and therefore missing from curated complex-level databases. In such cases, gene-level annotations still provide important functional context and reveal well-defined biological roles. Annotated sequence features—such as transmembrane domains—can be directly highlighted on the structure in the Mol* viewer (Fig. 5a–b).

**Figure 5.**
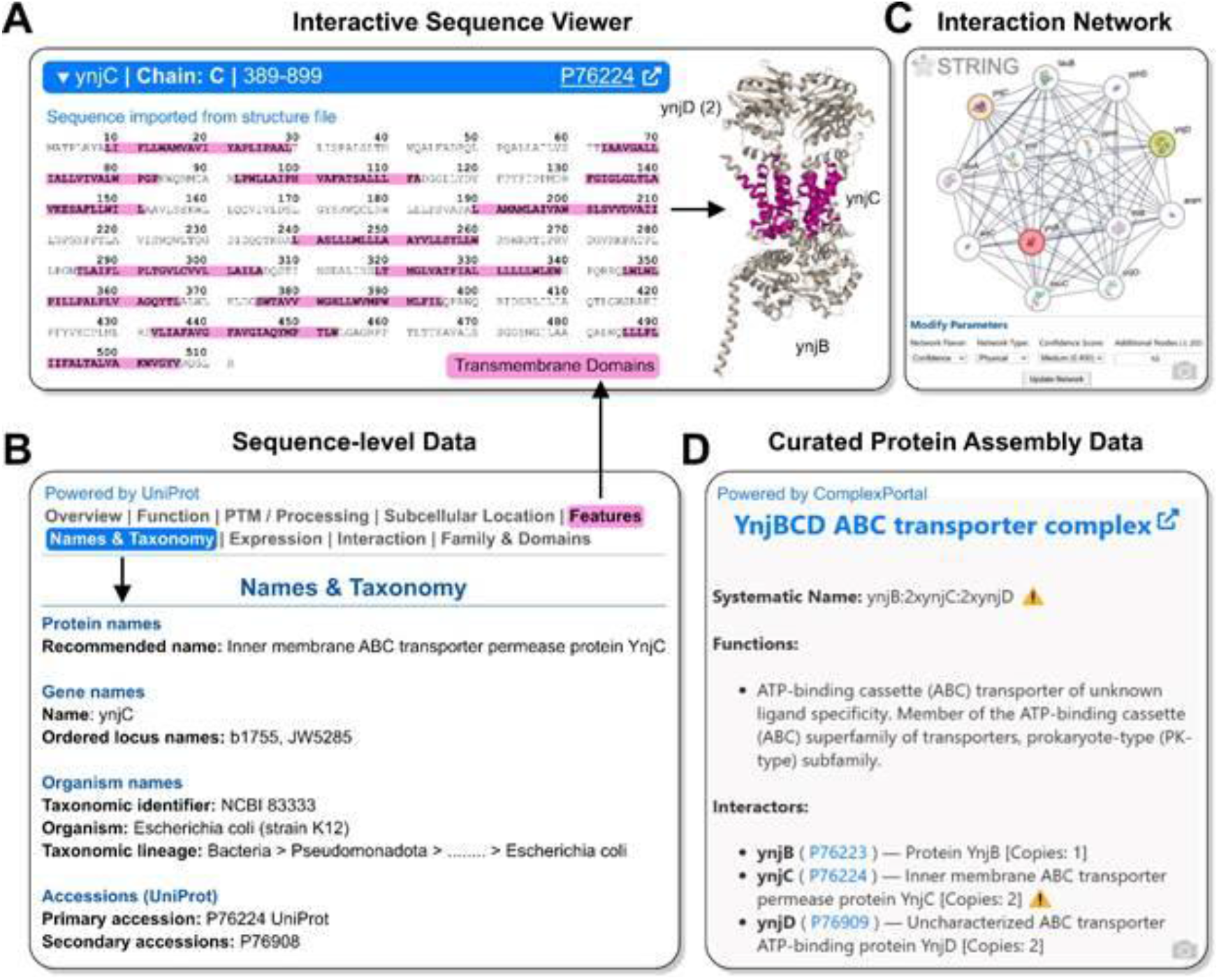
Integrating Biological Context into Structural Models. **(A)** The sequence viewer highlights selected residues on the 3D structure. **(B)** UniProt sequence features are displayed and linked to the structure viewer. **(C)** A STRING interaction network visualizes evidence of physical association among modeled subunits. **(D)** ComplexPortal annotations are shown for matching assemblies, including subunit composition and functional descriptions.

At the complex level, QSProteome integrates data from ComplexPortal, displaying complex names, functions, and biological roles for models that match or overlap with known assemblies (Fig. 5d). If no direct match is found, QSProteome generates a composite summary from the UniProt annotations of individual subunits.

To assess protein–protein interactions, each model page includes an interactive STRING network (Fig. 5c). This visualization combines experimental evidence and computational predictions to evaluate whether the modeled subunits are likely to form a biologically relevant complex. Users can examine connection confidence, supporting evidence, and interaction types directly within the embedded viewer.

By embedding rich biological annotations alongside structural models, QSProteome ensures that users can evaluate both structural integrity and biological relevance. This layered context supports expert interpretation and broadens accessibility to non-specialists seeking functionally meaningful models.

### 5. A systematic approach for accelerating community-driven modeling efforts

While tools like AlphaFold and ColabFold have expanded access to structure prediction, large-scale modeling of quaternary protein complexes remains constrained by computational demands and the lack of coordinated infrastructure. In particular, the absence of a centralized pipeline for managing unmodeled complexes slows the completion of full structural proteomes. Addressing this gap requires a system that supports efficient, non-redundant, and community-scalable modeling.

QSProteome meets this need with a server-side distribution system that organizes and prioritizes missing protein complexes for modeling. For each unmodeled complex, QSProteome generates a ready-to-use AlphaFold 3 input file, available for download from the search results table. Users can launch individual modeling jobs with minimal setup. For batch efforts, the Repository Page offers downloadable JSON files containing up to 30 curated input jobs, formatted for direct submission to the AlphaFold server (Fig. 2c).

To avoid duplication and optimize use of computational resources, QSProteome employs a queuing system that distributes modeling tasks systematically. Each job is assigned to a user for a fixed period (e.g., 48 hours), during which others are temporarily restricted from claiming the same task. If no submission is received, the task is opened for reassignment. This approach ensures that modeling efforts are non-redundant, contributions are properly credited, and valuable compute time is not wasted.

By streamlining and distributing modeling workloads across the community, QSProteome accelerates the expansion of structural proteomes, particularly for complexes whose gene-stoichiometries are annotated in public databases (e.g., BioCyc and ComplexPortal) and in previously published large-scale modeling efforts^12,20-21^. To showcase QSProteome’s distributed queuing system, we converted the published gene-stoichiometries across these datasets into ready-to-use AlphaFold 3 input files and made them available through the QSProteome interface, enabling non-redundant community-driven modeling of more than 35,000 unique protein assemblies in under 14 weeks. Furthermore, the system is designed to easily incorporate additional targets, enabling quick onboarding of new datasets. Over time, failed or low-confidence predictions will help pinpoint current algorithmic limitations, guiding future improvements in modeling tools. Additionally, submitted models that partially overlap with known assemblies may support stoichiometry reannotation, improving the biological accuracy of existing databases (see Fig. S1).

### 6. A gamified workflow for large-scale model re-curation, demonstrated on ABC transporters

ATP-binding cassette (ABC) transporters are a ubiquitous superfamily of proteins that mediate the ATP-dependent translocation of a wide variety of substrates across membranes, playing essential roles in nutrient uptake, drug resistance, and cell signaling across all domains of life^29^.

Despite their biological importance, most ABC transporters lack complete quaternary structural models, and gene-level stoichiometry remains unresolved for the majority of species outside well-characterized model organisms.

To demonstrate QSProteome’s capacity for structured, community-driven refinement of quaternary assemblies, we developed the “ABC Game” (QSProteome.org/abc-game)—an interactive tool that guides users through recognition of ABC transporter architectures, identification of missing subunits, and classification of transporter families (Types I–VII). The game presents models of increasing complexity, asking players to inspect 3D structures, select any missing domains (e.g., ATP-binding cassette, transmembrane segments, substrate-binding units), and assign each complex to its correct family (Fig. 6a).

**Figure 6.**
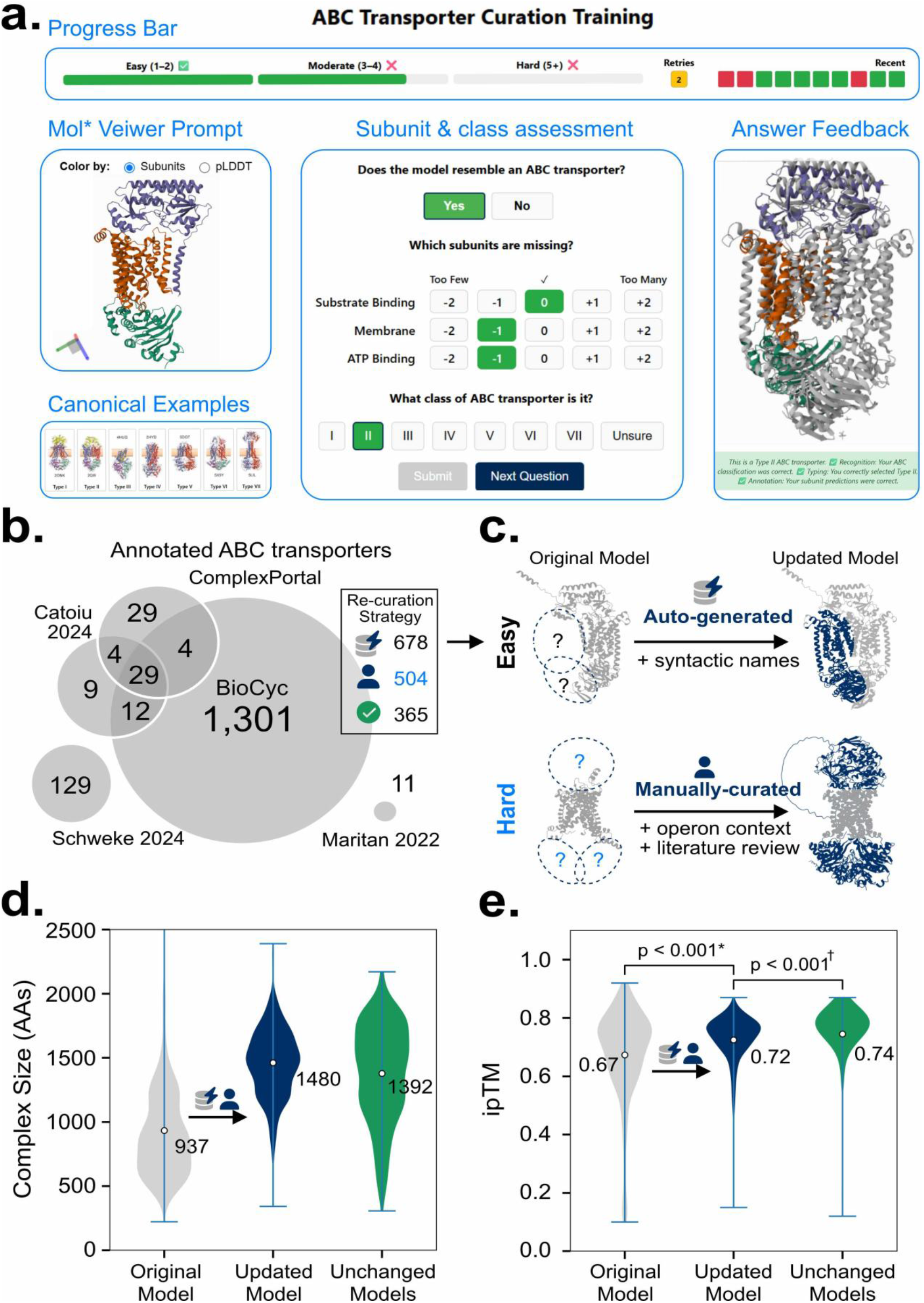
Model-guided refinement of ABC transporter assemblies through gamified annotation and automated curation. **(A)** The ABC Transporter Game guides users through model assessment tasks in a structured and engaging format. A progress bar tracks difficulty and performance (top), while a Mol* viewer (left) allows users to interact with the prompted model. Users assess whether the model resembles an ABC transporter and annotates subunit composition and class (center). Canonical examples provide visual reference for transporter types (bottom left), and feedback (including structural alignment to the correct canonical structure) is displayed after answer submission (right), reinforcing accurate identification and classification. **(B)** Initial ABC models were generated based on gene stoichiometries from public databases. The visual inspection and gene content of each model informed its classification for either automated or manual re-curation. **(C)** Minor stoichiometric issues were corrected via name parsing; major domain omissions triggered manual updates using operon context. **(D)** Amino acid length increased after curation, reflecting restoration of missing domains. **(E)** Inter-chain TM-scores (ipTM) improved post-curation, approaching confidence levels of known high-quality assemblies. (*paired Wilcoxon, ^†^Mann–Whitney)

Ten users completed the ABC Game and subsequently evaluated 1,547 unique ABC transporter models submitted to QSProteome (Table S1). These models originated from pre-annotated ABC assemblies in BioCyc, ComplexPortal, and previous large-scale modeling efforts. Based on visual inspection: 365 models (23.6%) were deemed correct and required no changes, 678 models (43.8%) had minor stoichiometric issues (e.g., a single permease domain where two are expected), and 504 models (32.6 %) were missing one or more essential domains (e.g., ATP-binding or substrate-binding subunits), representing “hard” cases (Fig. 6b).

For “easy” cases, QSProteome’s backend automatically parsed syntactic gene-product names, updated stoichiometries to reflect the visual assessment, and generated revised AlphaFold 3 input files (Fig. 6c, top). “Hard” cases underwent manual curation: we inspected operon context provided by BioCyc^30^, identified missing gene locus tags, and assigned copy numbers consistent with known operon structures and game annotations (Fig. 6c, bottom).

Recuration efforts uncovered 714 missing subunit types and 1,248 total corrections to subunit stoichiometry. After re-curation, the average amino acid count in each model increased from 937 to 1,480 residues (Fig. 6d), and inter-chain TM-scores (ipTM) improved from 0.67 to 0.72 (paired Wilcoxon *p* < 0.001). These post-curation scores matched those of well-characterized ABC complexes (Mann–Whitney *p* < 0.001; Fig. 6e). Overall, this gamified, community-curated workflow restored missing subunits and improved quaternary structure across 1,547 ABC complexes—a scalable framework for systematic reannotation of oligomeric assemblies. A fully interactive before-and-after view of all curated complexes–displaying gene stoichiometries, ipTM scores, curation strategy, and cross-referenced databases—is available at: https://qsproteome.org/abc-final-curation.

This curated collection of ABC transporters represents the most comprehensive, structure-resolved resource available for this protein superfamily. Previous studies have used structural features to classify subsets of ABC transporters by domain architecture or substrate specificity^31-33^, but such efforts have been constrained by limited model availability and the time-intensive manual curation required to resolve stoichiometric inconsistencies. The re-curated QSProteome dataset overcomes these barriers, enabling large-scale, cross-species analyses of structure-function relationships, phylogenetic groupings, and substrate prediction. Foldseek alignments^34^ reveal recurring quaternary architectures across species (Fig. S3), laying the groundwork for future integrative studies. This scalable reannotation framework can be extended to other large, functionally diverse protein families with complex quaternary architectures–such as two-component systems^35^, chaperonins^36^, secretion systems^37^ and LSm proteins^38^–where stoichiometric ambiguity and structural heterogeneity play critical roles.

## Discussion

QSProteome represents the first interactive repository for community-sourced, computationally predicted higher-order protein complexes. Unlike static structure databases, QSProteome provides real-time access to high-confidence models alongside tools for exploration, validation, and annotation. The platform enables researchers to instantly assess model quality while benefiting from thousands of structures that have already been computed and vetted by the community. By supporting user uploads, prioritized modeling queues, and seamless integration with curated biological databases, QSProteome lays the groundwork for continuous and scalable modeling across diverse organisms and biological systems.

A central feature of QSProteome is its comprehensive, transparent framework for model validation, designed to bring CASP-like benchmarking^39^ to scale. All models are aligned to related experimental structures using MM-align, while the conservation of quaternary shape in homo-oligomeric models is evaluated with QSalign. These orthogonal comparisons are visualized at both the model (Fig. 2c & Fig. 4d) and dataset (Fig. S2) levels, enabling systematic assessment across tens of thousands of complexes. Furthermore, QSProteome supports the large-scale deployment of emerging scoring metrics, such as interface-focused chain-minimum PAE^16^ and ipSAE^28^, when they are derivable from standard model outputs. In contrast, scoring methods that depend on intermediate AlphaFold outputs cannot be applied retrospectively (e.g., residue–pair error probability distributions used by actifpTM^40^). To maximize benchmarking, future prediction engines should be designed with emerging scoring metrics in mind, exposing intermediate data to enable retrospective application of new metrics across existing model libraries. By enabling large-scale deployment of both conventional and emerging scoring metrics across a broad structural landscape, QSProteome provides valuable insights for model developers, CASP assessors, and biologists alike—helping to identify failure modes, guide improvements in prediction algorithms, and highlight promising targets for experimental follow-up.

Beyond structural accuracy, QSProteome enhances biological interpretability by integrating contextual data from UniProt, STRING, and ComplexPortal. This multi-layered annotation framework links each structural model to relevant sequence features, interaction networks, and curated complex definitions. The result is a resource that supports both structural and functional analysis, making predicted models more accessible to a broader scientific community. By embedding structure predictions within well-established biological knowledge, QSProteome helps translate computational output into meaningful biological insight.

QSProteome also establishes a robust framework for organized, large-scale community modeling. The Upload Portal allows researchers to contribute models seamlessly, while a structured queuing system ensures efficient and non-redundant expansion of the database. Since the initial drafting of this work, QSProteome now hosts models for nearly all protein complexes annotated in BioCyc and ComplexPortal, demonstrating the platform’s scalability and broad coverage. Because QSProteome is directly integrated with these external databases, any updates—such as newly curated complexes or modified gene stoichiometries—are automatically incorporated into the modeling pipeline. Moreover, the platform’s architecture is highly extensible: it can readily accommodate new sources of protein assemblies, including gene stoichiometry datasets from previous large-scale modeling efforts^12,20-21^. This flexibility ensures that QSProteome can evolve in parallel with the rapidly expanding landscape of structural systems biology.

Predicted structural models can also offer valuable insights for refining curated biological knowledge, particularly when they suggest unexpected subunit composition or stoichiometry (e.g., Fig. S1). However, case-by-case expert evaluation is difficult to scale. Gamified platforms such as *Foldit* and *Borderlands Science* have shown that scientific discovery can be streamlined by engaging users in structured annotation tasks^41-42^. Building on this idea, we developed a lightweight, guided curation workflow within QSProteome, focused on ABC transporters. By combining AlphaFold predictions with game-like feedback and visual templates, we enabled 10 users to rapidly re-curate 1,547 ABC transporter complexes, demonstrating that predicted models can drive efficient, high-throughput refinement of biological databases with accuracies comparable to previously well-characterized systems (e.g., Fig. 6e). Together, these results underscore QSProteome’s role as a dynamic bridge between structural prediction and expert curation.

QSProteome’s cloud-based architecture is designed to scale with advances in modeling, enabling retrospective assessment and the identification of candidates for re-computation or reannotation. Low-confidence predictions can highlight systematic weaknesses in current methods, while high-confidence models—particularly those that differ from known assemblies—may inform updates to misannotated complexes. To promote reverse-interoperability, we have begun exposing public APIs for retrieval of QSProteome models, data and embeddable Mol* widgets corresponding to any BioCyc, ComplexPortal, UniProt, and PDB entry, available at qsproteome.org/api-usage. Looking forward, we hope to enable interactive widgets that allow users to submit stoichiometry refinements directly from partner websites and pass them into our distributed modeling queue to produce AlphaFold-3–ready input files, thereby harnessing external user communities to accelerate curation and extend quaternary structural proteome coverage. Crucially, by embedding predicted complexes within a structured comparative framework, QSProteome serves not merely as a protein archive, but as a platform for continuous evaluation, contextualization, and improvement of modeled structures.

In doing so, QSProteome also lays the groundwork for broader applications: beyond validation and annotation, it can become a valuable resource for whole-cell 3D reconstructions^21^ and genome-scale models of macromolecular expression^43-45^, where subunit-level oligomeric assemblies are essential building blocks. By providing these “recipes”—annotated, validated, and biologically grounded—QSProteome empowers researchers to incorporate realistic molecular components into cellular-scale simulations and visualizations.

The success of community-led platforms like Global Natural Products Social Molecular Networking^46^ demonstrates the transformative potential of socializing biological data–turning raw experimental outputs into dynamic, collaborative resources. Inspired by this model, QSproteome aims to foster a similar culture of open sharing, iterative refinement, and cross-disciplinary collaboration in the domain of modeled protein complexes. We invite the broader scientific community—across disciplines and expertise levels—to engage with and contribute to QSProteome. Community contributions—whether uploads, refinements, benchmarking, or suggestions for new large-scale projects for modeling—advance our shared goal of understanding the molecular machinery of life.

## Data Availability

QSProteome is freely available online for academic and non-profit use at https://QSProteome.org.

All coordinate files, metrics files, and configuration files are available for download. Model pages can be accessed at https://QSProteome.org/entry/{QS-entryId}. We encourage community members to submit their own modeling results directly on our Upload Portal. Supplementary data is provided in interactive form at https://QSProteome.org/abc-final-curation and https://QSProteome.org/abc-foldseek. The ABC game is available at https://QSProteome.org/abc-game. For technical methods, refer to the Supplementary Materials.

## Acknowledgements

We thank Peter Karp and Suzanne Paley for their guidance and support with BioCyc data access and integration. We also acknowledge the contributions of Shadee Hemaidan, Tanya Kumar, and Maahi Shah for their dedicated efforts in data gathering, annotation and beta-testing of the site. Author Contributions: E.A.C. conceptualized the project, led project development, performed analyses, implemented the backend and frontend, and drafted the manuscript. D.K., B.R., K.V., Z.W., J.A., J.S., R.A., M.H., K.A., A.T., I.G., R.A., C.F., J.N., B.T., and L.O.M. contributed to data generation, annotation, and beta testing. Z.W. participated in project discussions, contributed frontend code, and provided insights on transporter gamification. B.O.P. provided mentorship, supervision, conceptualization, financing, facilities, and overall project guidance. All authors contributed to manuscript editing and review.

## Ethics Declarations

The authors declare no conflicts of interest.

## References

1. Jumper, J., Evans, R., Pritzel, A. et al. Highly accurate protein structure prediction with AlphaFold. Nature. 596, 583–589 (2021). 10.1038/s41586-021-03819-2

2. Mirdita, M., Schütze, K., Moriwaki, Y. et al. ColabFold: making protein folding accessible to all. Nat. Methods 19, 679–682 (2022). 10.1038/s41592-022-01488-1

3. Lupas A., Pereira J., Alva V., et al. The breakthrough in protein structure prediction. Biochem. J. 478(10):1885–1890 (2021). 10.1042/BCJ20200963

4. Kryshtafovych A., Schwede T., Topf M., et al. Critical assessment of methods of protein structure prediction (CASP)—Round XIV. Proteins. 89(12):1607–1617 (2021). 10.1002/prot.26237

5. Evans R., O’Neil M., Pritzel A., et al. Protein complex prediction with AlphaFold-Multimer. bioRxiv. (2022). 10.1101/2021.10.04.463034

6. Tauriello, G., Waterhouse A,. Haas J., et al. ModelArchive: A Deposition Database for Computational Macromolecular Structural Models. J. Mol. Biol. 437(15):168996 (2025) 10.1016/j.jmb.2025.168996

7. Varadi, M., Bertoni, D., Magana, P., et al. AlphaFold Protein Structure Database in 2024: Providing structure coverage for over 214 million protein sequences. Nucleic Acids Res. 52(D1), D368–D375 (2023). 10.1093/nar/gkad1011

8. Dey, S., Ritchie, D. & Levy, E. PDB-wide identification of biological assemblies from conserved quaternary structure geometry. Nat. Methods. 15, 67–72 (2018). 10.1038/nmeth.4510

9. Dey S., Levy E.D. PDB-wide identification of physiological hetero-oligomeric assemblies based on conserved quaternary structure geometry. Structure. 29(11):1303–1311 (2021). 10.1016/j.str.2021.07.012.

10. Dey S, Prilusky J, Levy ED. QSalignWeb: A Server to Predict and Analyze Protein Quaternary Structure. Front Mol Biosci. 8:787510 (2021). 10.3389/fmolb.2021.787510

11. Schweke H., Xu Q,. Tauriello G., et al. Discriminating physiological from non-physiological interfaces in structures of protein complexes: a community-wide study. Proteomics. 23(17):e2200323 (2023). 10.1002/pmic.202200323

12. Schweke, H., Pacesa M., Levin T., et al. An atlas of protein homo-oligomerization across domains of life. Cell. 187(4):999–1010 (2024). 10.1016/j.cell.2024.01.022

13. The UniProt Consortium. UniProt: the Universal Protein Knowledgebase in 2025. Nucleic Acids Res. 53(D1):D609–D617 (2025). 10.1093/nar/gkae1010

14. Meldal B.H.M., Perfetto L., Combe C., et al. Complex Portal 2022: new curation frontiers. Nucleic Acids Res. 50(D1):D578–D586 (2022). 10.1093/nar/gkab991

15. Szklarczyk D., Kirsch R., Koutrouli M., et al. The STRING database in 2023: protein-protein association networks and functional enrichment analyses for any sequenced genome of interest. Nucleic Acids Res. 51(D1):D638–D646 (2023). 10.1093/nar/gkac1000

16. Abramson, J., Adler, J., Dunger, J. et al. Accurate structure prediction of biomolecular interactions with AlphaFold 3. Nature. 630, 493–500 (2024). 10.1038/s41586-024-07487-w

17. Karp, P.D., Billington, R., Caspi, R., et al. The BioCyc collection of microbial genomes and metabolic pathways. Brief Bioinform. 20(4):1085–1093 (2019). 10.1093/bib/bbx085

18. Karp, P.D., Paley, S., Caspi, R., et al. The EcoCyc database in 2023. EcoSal Plus. 11(1):eesp00022023 (2023). 10.1128/ecosalplus.ESP-0002-2023

19. Caspi, R., Billington, R., Keseler, I.M., et al. The MetaCyc database of metabolic pathways and enzymes - a 2019 update. Nucleic Acids Res. 48(D1):D445–D453 (2020). 10.1093/nar/gkz862

20. Catoiu E.A., Mih N., Lu M., Palsson B.O. Establishing comprehensive quaternary structural proteomes from genome sequence. eLife. 13:RP100485 (2024). 10.7554/eLife.100485.1

21. Maritan M., Autin L., Karr J., Covert M.W., Olson A.J., Goodsell D.S. Building structural models of a whole mycoplasma cell. J. Mol. Biol. 434(2):167351 (2022). 10.1016/j.jmb.2021.167351

22. Ahmad, S., da Costa Gonzales L.J., Bowler-Barnett, E.H., et al. The UniProt website API: facilitating programmatic access to protein knowledge Nucleic Acids Res. 53(W1):W547–W553 (2025). 10.1093/nar/gkaf394.

23. Updated resources for exploring experimentally-determined PDB structures and Computed Structure Models at the RCSB Protein Data Bank. Nucleic Acids Res. 53(D1):D564–D574 (2025). 10.1093/nar/gkae1091

24. Mukherjee, S. & Zhang Y. MM-align: a quick algorithm for aligning multiple-chain protein complex structures using iterative dynamic programmin. Nucleic Acids Res. 37(11), e83 (2009). 10.1093/nar/gkp318

25. O’Leary, N.A., Cox, E., Holmes, J.B., et al Exploring and retrieving sequence and metadata for species across the tree of life with NCBI Datasets. Sci Data. 11, 732 (2024). 10.1038/s41597-024-03571-y

26. Hastings J., Owen G., Dekker A., et al. ChEBI in 2016: Improved services and an expanding collection of metabolites. Nucleic Acids Res. 44(D1):D1214–D1219 (2016). 10.1093/nar/gkv1031

27. Sehnal, D., Bittrich S., Deshpande M., et al. Mol* Viewer: modern web app for 3D visualization and analysis of large biomolecular structures. Nucleic Acids Res. 49(W1):W431–W437 (2021). 10.1093/nar/gkab314

28. Dubrack R.L. Res ipSAE loquunt: What’s wrong with AlphaFold’s ipTM score and how to fix it. bioRxiv. 2025. doi: 10.1101/2025.02.10.637595

29. Rees, D., Johnson, E. & Lewinson, O. ABC transporters: the power to change. Nat. Rev. Mol. Cell Biol. 10, 218–227 (2009). 10.1038/nrm2646

30. Herson, J., Krummenacker, M., Spaulding, A., et al. The Genome Explorer genome browser. mSystems. 9(7):e0026724 (2024). 10.1128/msystems.

31. Mahendran, A. & Orlando, B. J. Genome wide structural prediction of ABC transporter systems in Bacillus subtilis. Front. Microbiol. 15, 1469915 (2024). 10.3389/fmicb.2024.1469915

32. Hu, Y., Guo, Y., Li, M., et al. A consensus subunit-specific model for annotation of substrate specificity for ABC transporters. RSC Adv. 52, 42009–42019 (2015). 10.1039/C5RA05304H

33. Berntsson, R.P.-A., Smits, S.H.J., Schmitt, L., et al. A structural classification of substrate-binding proteins. FEBS Lett. 584(12):2606–2617 (2010). 10.1016/j.febslet.2010.04.043

34. van Kempen, M., Kim, S.S., Tumescheit, C., et al. Fast and accurate protein structure search with FoldSeek. Nat. Biotechnol. 42, 243–246 (2024). 10.1038/s41587-023-01773-0

35. Edkins, A.L., Boshoff, A. General Structural and Functional Features of Molecular Chaperones. In: Shonhai, A., Picard, D., Blatch, G.L. (eds) Heat Shock Proteins of Malaria. Advances in Experimental Medicine and Biology, vol 1340. Springer, Cham. 10.1007/978-3-030-78397-6_2

36. Gao R, Stock AM. Biological insights from structures of two-component proteins. Annu. Rev. Microbiol. 63, 133–54 (2009). 10.1146/annurev.micro.091208.073214.

37. Costa, T.R.D., Patkowski, J.B., Macé, K. et al. Structural and functional diversity of type IV secretion systems. Nat. Rev. Microbiol. 22, 170–185 (2024). 10.1038/s41579-023-00974-3

38. Mura, C., Randolph, P.S., Patterson, J., & Cozen, A.E. Archaeal and eukaryotic homologs of Hfq: A structural and evolutionary perspective on Sm function. RNA Biol. 10(4):636–51 (2013). doi: 10.4161/rna.24538.

39. Kryshtafovych, A., Schwede, T., Topf, M., Fidelis,. K, & Moult, J. Critical assessment of methods of protein structure prediction (CASP)—Round XV. Proteins. 91(12): 1539–1549 (2023). doi:10.1002/prot.26617

40. Varga, J.K., Ovchinnikov, S., & Schueler-Furman, O. actifpTM: a refined confidence metric of AlphaFold2 predictions involving flexible regions. Bioinformatics. 41(3):btaf107 (2025). doi: 10.1093/bioinformatics/btaf107

41. Kleffner, R., Flatten, J., Leaver-Fay, A., et al. Foldit Standalone: a video game-derived protein structure manipulation interface using Rosetta, Bioinformatics. 33(17) 2765–2767 (2017). 10.1093/bioinformatics/btx283

42. Sarrazin-Gendron, R., Ghasemloo Gheidari, P., Butyaev, A. et al. Improving microbial phylogeny with citizen science within a mass-market video game. Nat. Biotechnol. 43, 76–84 (2025). 10.1038/s41587-024-02175-6

43. Lloyd C.J., Ebrahim A., Yang L., et al. COBRAme: A computational framework for genome-scale models of metabolism and gene expression. PLoS. Comput. Biol. 14(7):e1006302 (2018). 10.1371/journal.pcbi.1006302

44. Chen, K., Gao, Y., Mih, N., et al. Thermosensitivity of growth is determined by chaperone-mediated proteome reallocation. PNAS. 114(43):11549–11553 (2017). 10.1073/pnas.1705524114

45. Zhao, J., Chen, K., Palsson, B.O., & Yang, L. StressME: Unified computing framework of Escherichia coli metabolism, gene expression, and stress responses. PLoS. Comput. Biol. 20(2):e1011865 (2024). 10.1371/journal.pcbi.1011865

46. Wang, M., Carver, J., Phelan, V. et al. Sharing and community curation of mass spectrometry data with Global Natural Products Social Molecular Networking. Nat Biotechnol. 34, 828–837 (2016). 10.1038/nbt.3597

