## Supplemental Material for "QSProteome: A Community-Driven Interactive Platform for Large-Scale Exploration and Evaluation of Predicted Protein Complex Structures"

**This document contains:**

Figure S1-S3
Table S1
Supplementary Methods

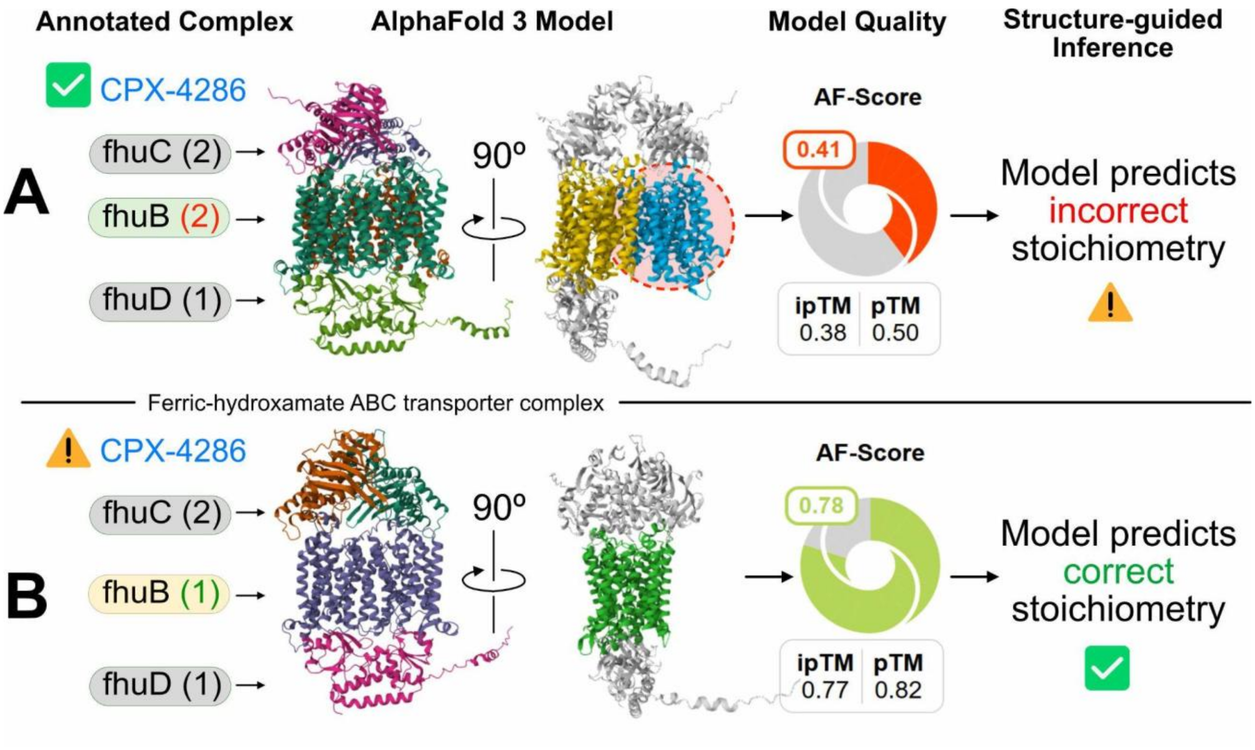


**Figure S1. Structure-guided correction of annotated stoichiometry in ComplexPortal.** Two AlphaFold 3 models of the ferric-hydroxamate ABC transporter complex (ComplexPortal: CPX-4286) illustrate how high-confidence predictions can challenge or refine reference annotations. **(A)** The ComplexPortal entry specifies two copies of *fhuB*, but the AlphaFold 3 produces a low confidence model (AF-score = 0.41; ipTM = 0.38), suggesting incorrect stoichiometry. **(B)** A higher-confidence model of the same complex (AF-score = 0.78; ipTM = 0.77) predicts one copy of *fhuB*, consistent with the structural assembly and improved model quality. This example supports the use of structure predictions to guide the curation of stoichiometric annotations in protein complex databases.

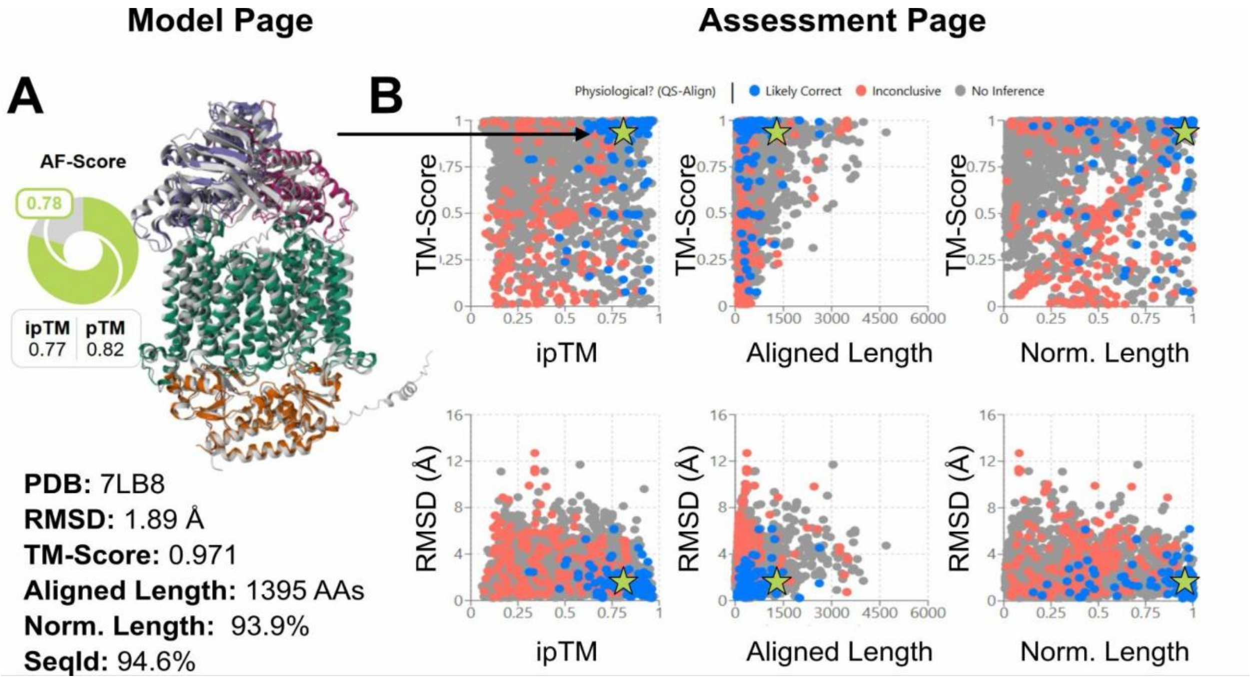


**Figure S2. Model-level and global structural alignment assessments using MM-align. (A)** The model page for an AlphaFold 3 prediction displays the structural alignment results against experimentally determined complexes in the PDB. Key statistics—RMSD (1.89 Å), TM-score (0.971), aligned length (1395 amino acids), normalized alignment length (93.9%), and sequence identity (94.6%)—are shown alongside AlphaFold-derived confidence scores (ipTM, pTM). In the Mol* viewer, the superimposed model (white) and aligned PDB structure (colored) enable direct visual comparison. **(B)** The Assessment Page provides a global summary of MM-align results across all models. Each plot visualizes TM-score or RMSD against metrics such as ipTM, aligned length, or normalized alignment length. Points are color-coded by validation status: blue indicates QS-align-supported models (likely correct), red indicates inconclusive results, and gray represents models without structural inference. The green star highlights the currently viewed model. On hover, model images and alignment metrics are displayed; clicking a point navigates to the corresponding model page. The model shown is the same as in Figure S1B.

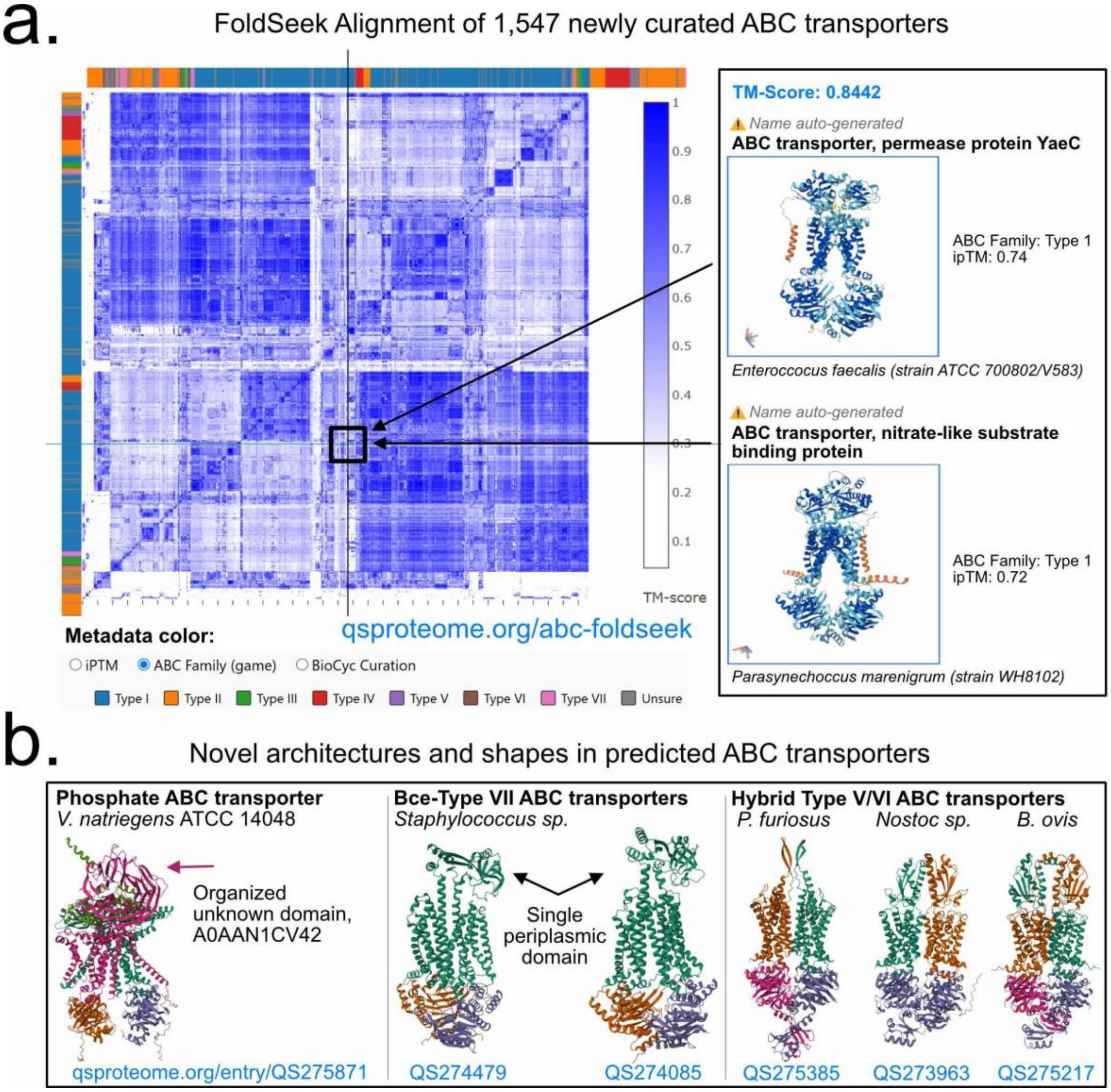


**Figure S3. Foldseek-based structural alignment of 1,547 curated ABC transporter complexes. (A)** A pairwise TM-score similarity matrix reveals recurring quaternary architectures among curated ABC transporter models. Colored bars along the matrix axes indicate predicted ABC family (Type I–VII) from the ABC Game. Representative structures from a high-similarity cluster illustrate conserved architecture across distant taxa. This structural clustering highlights shared assembly principles and provides a foundation for future large-scale classification of transporter families. Full interactive matrix with additional coloring schemes (BioCyc overlap & ipTM) is available at qsproteome.org/abc-foldseek. **(B)** Interesting non-canonical quaternary shapes identified from the alignment matrix.

**Table S1. Community-driven visual assessment of ABC transporter assemblies via the ABC Game.** Ten users participated in the interactive ABC Game and provided visual assessments of ABC transporters uploaded to QSProteome. Metrics include the number of training responses* and total training time (in minutes)† prior to live deployment, as well as live annotations‡ and consensus-based accuracy§. These crowd-sourced annotations directly informed the re-curation and remodeling of 1,547 unique ABC transporter assemblies.

| User | Training Responses* | Training Time (minutes)† | Live Annotations‡ | Live Accuracy§ |
|---|---|---|---|---|
| A | 207 | 241 | 59 | 94.9% |
| B | 164 | 31 | 2189 | 90.3% |
| C | 176 | 139 | 650 | 86.6% |
| D | 279 | 206 | 409 | 86.6% |
| E | 411 | 197 | 2189 | 86.2% |
| F | 648 | 134 | 972 | 80.9% |
| G | 313 | 198 | 605 | 80.3% |
| H | 515 | 189 | 2113 | 75.6% |
| I | 468 | 275 | 130 | 75.4% |
| J | 374 | 125 | 90 | 73.3% |

*Training metrics vary by user due to differences in game versions and evolving difficulty thresholds during development.

†Calculated as the cumulative time between consecutive training responses, each separated by less than 3 minutes.

‡Multiple initial models processed through the re-curation pipeline may correspond to the same final stoichiometry.

§Accuracy reflects consensus agreement, where an annotation is considered correct if more than 50% of subunit assessments for a model match across users.

### Supplementary Methods

#### Model Upload and Processing

Users submit models through the QSProteome Upload Portal (https://qsproteome.org/upload), which accepts outputs from the AlphaFold 3 webserver or legacy ColabFold. Each uploaded folder is parsed to identify coordinate files (CIF or PDB), metrics files (scores.json / summary_confidences.json / full_data.json), and configuration files (config.json). A SHA-256 hash of the coordinate file is computed and saved in a relational database. Redundant uploads are avoided by cross-checking newly computed hashes against previously stored entries. The best-ranked model from each folder is selected for downstream analysis based on internal AlphaFold confidence metrics.

#### Model Evaluation and Structural Alignment

Uploaded models are evaluated using AlphaFold-derived confidence scores: inter-protein TM-score (ipTM), predicted TM-score (pTM), predicted Local Distance Difference Test (pLDDT), and Predicted Aligned Error (PAE). Homo-oligomeric models are automatically submitted to QSalignWeb (https://shmoo.weizmann.ac.il/qsab/main) [ref #8] , which assesses evolutionary conservation of quaternary shape. Submissions are performed through automated browser-based interaction using Selenium WebDriver, which replicates manual uploads in a headless Chrome session. Upon successful submission, QSProteome periodically polls the QSalignWeb server for alignment results, which are then downloaded and parsed for integration. Related PDB structures are identified via UniParc and UniProt cross-references based on subunit-level sequence similarity. These structures are downloaded from RCSB PDB and locally aligned to the uploaded model using MM-align [ref #20], a command-line tool for structural comparison of multi-chain protein complexes. We used the following command: “MMalign <ref.pdb> <model.pdb> -o <outdir> -full T -ter 1”, where -full T enables full-atom TM-score calculation and -ter 1 enforces terminal atom alignment. MM-align outputs key alignment statistics including root-mean-square deviation (RMSD), TM-score, aligned length, and sequence identity. The top-ranked structural alignment and the corresponding superimposed coordinate file are displayed on each model page for direct visual comparison. Aggregate alignment metrics are displayed at qsproteome.org/summary. On the browser-side, we retrospectively calculate and display the chain-minimum PAE score (Abramson, 2024) and the ipSAE score (Dunbrack, 2025) for each model.

#### Biological Annotation and Integration

Each model is annotated using UniProt (gene names, protein names, ligands, functions), ComplexPortal (stoichiometry and biological role), and STRING (subunit interaction networks). These annotations are displayed alongside structural visualizations and dynamically updated via REST API calls. When metadata is displayed on the Model and Search Pages, all cross-referenced databases (PubMed, ChEBI [ref #31], ComplexPortal, UniProt, NCBI) are hyperlinked to external pages for direct navigation.

#### Model Categorization and Repository Search

Each uploaded model is categorized into one of three classes based on its subunit stoichiometry relative to its source of the model inputs (BioCyc complexes, ComplexPortal entry, previous modelling efforts [ref# 10, ref# 22, ref #23]): (1) exact match, where the gene composition and stoichiometry fully align; (2) partial overlap, where one or more subunits match but the stoichiometry differs; or (3) unmatched, where no corresponding source assembly exists.

All metadata—including gene names, complex IDs, protein functions, ligands, taxonomy, and AlphaFold confidence metrics—is indexed in the database to support full-text and faceted search. Users can perform keyword-based searches or use the Advanced Search interface for structured filtering by organism, gene composition, or model availability. Keyword searches implement a fuzzy matching algorithm to accommodate spelling variations and synonyms, while advanced filters enforce exact field-based matching or threshold cutoffs. Full search table results can be downloaded on the user-interface.

#### Organized Community Modeling Workflow

To enable large-scale modeling while minimizing redundancy, QSProteome implements a structured queuing system for distributing modeling tasks. The Repository Page displays all curated assemblies from different cross-referenced databases alongside their current modeling status in QSProteome. Only exact gene-stoichiometric matches to input complexes are shown.

When a user initiates a batch download, the backend assigns a curated set of 30 unmodeled input protein complexes. For each complex, an AlphaFold 3-compatible input file (in JSON format) is automatically generated. This file includes the parsed subunit stoichiometry and representative amino acid sequences corresponding to the biologically active isoform annotated in cross-referenced databases or publications. The input files are packaged into a single .json archive and reserved exclusively for the requesting user for a defined period (typically 48 hours). During this reservation window, the selected complexes are temporarily removed from the modeling queue to prevent duplication.

Upon model upload, QSProteome verifies the presence of all required files and updates the corresponding repository entry to reflect completion. If no model is submitted within the reservation window, the task is released back into the pool of available jobs. All modeling actions are logged in a PostgreSQL backend and coordinated through a Redis-backed task queue implemented using Bull.

This reservation-based queuing logic ensures efficient structural coverage, prioritizes unmodeled complexes, and enables distributed model generation across a growing contributor base.

#### System Infrastructure

QSProteome is hosted on AWS infrastructure, including: a lightweight 4GB EC2 instance for server and queue management, S3 for storage of uploaded models and metadata, RDS for relational data management. The system uses a Redis-backed queuing service to manage asynchronous tasks such as upload validation, QSalign submissions, annotation retrieval, and structural alignment.

#### Gamified visual assessment of ABC transporters

To facilitate community-guided re-curation of ABC transporter assemblies, we developed an interactive tool called the ABC Game, available at https://qsproteome.org/abc-game. The game is powered by a training dataset of 400 annotated ABC transporter models. Each model was labeled with subunit presence/absence scores (−2 to +2) for three ABC domains—substrate-binding, membrane-spanning, and ATP-binding—and manually assigned to one of seven subfamilies (Types I–VII) based on literature and structural inspection.

To quantify task complexity, each model was assigned a difficulty score calculated as follows: +1 for each incorrect subunit, +1 for uncertain subfamily classification, +2 for missing membrane-spanning domains, +2 if flagged as structurally "unusual". For example, a Type I transporter with an extra substrate-binding domain receives a score of 1, while an incomplete Type II lacking both permeases and family annotation would score 4. To increase model diversity, we added 400 non-ABC transporter models to the dataset, selected from search results using the query "transporter."

The ABC game engine progresses users through four sequential phases: ABC recognition, subunit annotation, subfamily classification, and a certification phase, in which prompts from phases 1-3 are combined. Each phase presents models in order of increasing difficulty: Easy (score 0–1), Moderate (2–3), and Hard (4+). To advance, users must achieve minimum rolling accuracies of 90%, 80%, and 70% over their last 10 responses at Easy, Moderate, and Hard levels, respectively. A retry mechanism reintroduces missed models after 4–5 additional submissions to reinforce learning.

Ten users completed certification and contributed annotations to the live database (models sourced from https://qsproteome.org/search?query=abc%20transporter). Their annotations showed consensus-based agreement rates ranging from 73% to 94% (Table S1).

#### Text-based identification of genes encoding missing ABC subunits.

To assign each gene product in the initial ABC transporter models to one of three functional categories—ATP-binding, membrane-spanning, or substrate-binding—we developed a keyword-weighting script to analyze UniProt metadata. For each gene, we parsed the associated UniProt entry and collected text fields including protein names, subcellular localization terms, topologies, similarity comments, cross-references, and feature descriptions. A set of curated, weighted keyword dictionaries was used to score matches across each subunit class. For example, the presence of "solute-binding" or "periplasmic binding protein" in the metadata strongly favored classification as a substrate-binding subunit, while keywords such as "permease" and "multi-pass membrane" indicated membrane-spanning subunits. ATP-binding subunits were inferred based on hits to phrases like "ATPase" and "ATP-binding". The subunit class with the highest cumulative score was assigned to each gene, and a confidence score was calculated as the winning class's weighted score divided by the total score. This classification provided the initial domain-level annotation used to build and evaluate candidate ABC transporter assemblies.

#### Operon-guided identification of genes encoding missing ABC subunits.

Text-based classification methods are limited to genes already present in the structural model and cannot detect missing components. In cases where domains were absent, we used the BioCyc genome browser to inspect operon context and retrieved AlphaFoldDB structures for candidate genes to evaluate quaternary assembly plausibility. This was especially critical for homo-dimeric models of ABC transporter domains derived from Schweke et al. (2024), which used QSAlign—a structure-based quality control pipeline restricted to homomeric assemblies. While these homodimers passed structural validation, they frequently consisted of two identical copies of a single membrane-spanning or ATP-binding subunit. Operon analysis revealed that the biologically correct complex should instead include two distinct, non-identical subunits—i.e., a heterodimeric architecture. We therefore replaced one copy of the homodimer with its paralogous partner and updated the stoichiometry accordingly. In addition to correcting misassigned homodimers, we used the same operon-guided approach to identify entirely missing components—such as substrate-binding proteins (SBPs) or ATPase subunits—and incorporated them into the reassembled complex to restore complete functional architecture.

#### All vs all foldseek alignment of ABC transporters

To assess structural redundancy within the curated collection of ABC transporter complexes, we performed an all-versus-all structural similarity search using Foldseek (github.com/steineggerlab/foldseek). A total of 1,547 oligomeric models were obtained from QSProteome, each representing a manually curated and re-modeled ABC transporter complex. These models were first flattened to single-chain structures by concatenating all polypeptide chains from each complex into a continuous model, allowing for simplified pairwise comparisons.

The flattened structures were searched against themselves (all vs all) using the foldseek easy-search command with the following parameters: exhaustive-search=1 max-seqs=1000000, min-seq-id=0, min-aln-len=0 e=1000000 and greedy-best-hits=0. This configuration forced an exhaustive pairwise comparisons with no alignment length or sequence identity constraints. The clustered pairwise TM-scores (Fig. S3) represent the structural diversity across the 1,547 curated ABC transporter models. An interactive plot is available at qsproteome.org/abc-foldseek.

#### Software and Tools

QSProteome and the data presented in this manuscript would not be possible without the following APIs, tools, and software:

- Colabfold (Alphafold v2) / Alphafoldserver.com (Alphafold v3) - Structure prediction
- QSalignWeb - Quaternary structure validation
- MM-align - Structural alignment
- Mol* Viewer - Interactive 3D structure visualization
- Node.js / React - Frontend + backend stack
- PostgreSQL / Redis - Data storage and queuing
- Axios / REST APIs - Data retrieval from UniProt, UniParc, STRING, ComplexPortal, PDB, AlphafoldDB, BioCyc
- FoldSeek (https://github.com/steineggerlab/foldseek) - Fast ABC transporter alignment
- ipSAE.py (modified from https://github.com/DunbrackLab/IPSAE) – Interaction scoring
